# TNFα-induced metabolic reprogramming drives an intrinsic anti-viral state

**DOI:** 10.1101/2022.04.14.488171

**Authors:** Jessica Ciesla, Isreal Moreno, Joshua Munger

**Affiliations:** Department of Biochemistry and Biophysics, University of Rochester School of Medicine and Dentistry, Rochester, New York, USA

## Abstract

Cytokines induce an anti-viral state, yet many of the functional determinants responsible for limiting viral infection are poorly understood. Here, we find that TNFα induces significant metabolic remodeling that is critical for its anti-viral activity. Our data demonstrate that TNFα activates glycolysis through the induction of muscle-specific hexokinase (HK2). Further, we show that glycolysis is broadly important for TNFα-mediated anti-viral defense, as its inhibition attenuates TNFα’s ability to limit the replication of evolutionarily divergent viruses. Stable-isotope tracing revealed that TNFα-mediated glycolytic activation promotes the biosynthesis of UDP-sugars (essential precursors of protein glycosylation) and that inhibition of glycolysis prevents the accumulation of several glycosylated anti-viral proteins. Consistent with the importance of glucose-driven glycosylation, glycosyl-transferase inhibition also attenuated TNFα’s ability to promote the anti-viral cell state. Collectively, our data indicate that cytokine-mediated metabolic remodeling is an essential component of the anti-viral response.

## Introduction

Viruses are obligate intracellular parasites that cause substantial human morbidity and mortality. Newly emergent viruses can cause pandemics, for example, SARS-CoV-2, which in 2020 was the 3^rd^ leading cause of death in the U.S with ∼375,000 deaths^1^. In contrast, endemic viruses frequently present a lower, but constant burden to human health. Human Cytomegalovirus (HCMV), an endemic β-Herpesvirus, causes severe disease in various immunosuppressed populations, including the elderly, cancer patients receiving immunosuppressive chemotherapy, transplant recipients, and AIDS patients^2, 3^, and is also a major source of congenital birth defects, with 1 in 1000 babies born exhibiting symptoms such as microcephaly, deafness or retinitis^4^.

Prevention of virally-associated morbidity requires a strong innate immune response^5^. These responses are frequently initiated through viral antigen recognition by cellular pattern recognition receptors (PRRs) that activate signal transduction cascades, and ultimately trigger the production of anti-viral cytokines, including Tumor Necrosis Factor-alpha (TNFα), Types 1, 2, and 3 Interferons (IFNs), and interleukins (ILs)^5^. These cytokines are critical for recruiting tissue-resident and circulating immune cells to the site of infection^6, 7^, but importantly, they also induce an anti-viral state in uninfected bystander cells. TNFα, for example, inhibits the replication of a variety of evolutionarily diverse viruses including Vesicular Stomatitis virus (VSV), Encephalomyocarditis virus (EMCV), Herpes Simplex virus (HSV)^8^, HCMV^9^, and Hepatitis C virus (HCV)^10^. For many cytokine-induced signaling pathways, much of the upstream signaling network has been elucidated, e.g., cytokine-receptor binding ultimately resulting in transcription factor activation and expression of cytokine-associated genes^11-13^. However, much less is known about the cellular biology associated with the institution of the anti-viral state. For example, while several cytokine-induced genes have been found to be important for intrinsic immune defense^13^, how these genes modulate normal cellular physiology to limit infection is largely unclear.

Metabolic reprogramming has emerged as a central feature of the functional responses of professional immune cells. For example, glycolytic regulation is important for B cell function^14, 15^, and aspects of fatty acid and mitochondrial metabolism are critical for T cell activation and the maintenance of memory T cells^16, 17^. Similarly, granulocytes, monocytes, and macrophages have all been found to rely on aspects of glycolytic and glutaminolytic metabolism to differentiate, polarize, infiltrate infected tissues, and phagocytose infected cells^18-22^. While metabolic remodeling in professional innate and adaptive immune cells has emerged as a critical component of a successful immune response, surprisingly little is known about how cytokine signaling impacts the metabolism of non-professional bystander cells or the potential role that cytokine-induced metabolic modulation contributes to limiting viral infection.

Here, we apply metabolomic approaches to elucidate how TNFα modulates cellular metabolism to support its anti-viral activity. We find that TNFα induces HK2 to activate glycolysis, and that TNFα-activated glycolysis funnels carbon towards UDP-sugar biosynthesis. Restricting glycolysis largely blocks TNFα’s ability to limit the replication of HCMV and two betacoronaviruses, OC43 and SARS-CoV-2. This loss of anti-viral activity coincides with substantially decreased expression of several intrinsic anti-viral factors, several of which are glycosylated. Consistent with an important role for glycosylation, we find that inhibition of glycosyltransferases also inhibits TNFα’s anti-viral activity. Together, our data indicate that TNFα-induced glycolysis promotes UDP-sugar turnover to support glycosylation, which is required for the expression of intrinsic anti-viral factors and the induction of an anti-viral cellular state.

## Results

### TNFα induces glycolysis as part of a broadly altered metabolic state

To elucidate how TNFα treatment impacts cellular metabolism, we employed LC-MS/MS to analyze cellular metabolic pools in vehicle-treated non-transformed human foreskin fibroblasts (HFFs) versus those treated with TNFα (Supplementary Table 1). Principal-component analysis (PCA) of the resulting data discriminated between vehicle and TNFα-treated samples in the first principal component (Fig. 1a). Further, hierarchical clustering of this data segregated TNFα treated samples from control samples (Fig. 1b). Together, these data suggest that TNFα treatment induces a distinct metabolic state. The relative abundance of twelve metabolites were significantly altered by TNFα treatment (Fig. 1c, & Supplementary Table 1.1). Notably, the largest metabolite increase was in kynurenine abundance, which was ∼20-fold more abundant in TNFα-treated cells relative to controls (Fig. 1c). Kynurenine is part of the tryptophan catabolic pathway that supports NAD^+^ biosynthesis^23, 24^, however, in this study NAD^+^ pools were significantly decreased upon TNFα treatment (Fig. 1c). Intriguingly, a similar response, i.e., increased kynurenine and decreased NAD^+^ levels, was recently shown to occur after inflammatory challenge in macrophages, and subsequently found to be important for proper innate immune responses^25^. In addition, two of the most significantly increased metabolites, ribose-phosphate and sedoheptulose-7-phosphate, are part of the pentose phosphate pathway (Fig. 1c & d). Hexose-phosphate was also significantly increased by TNFα treatment (Fig. 1c & d), as was UDP-glucose, a key glycosylation precursor and glycogen building block (Fig. 1c & d).

**Figure 1.**
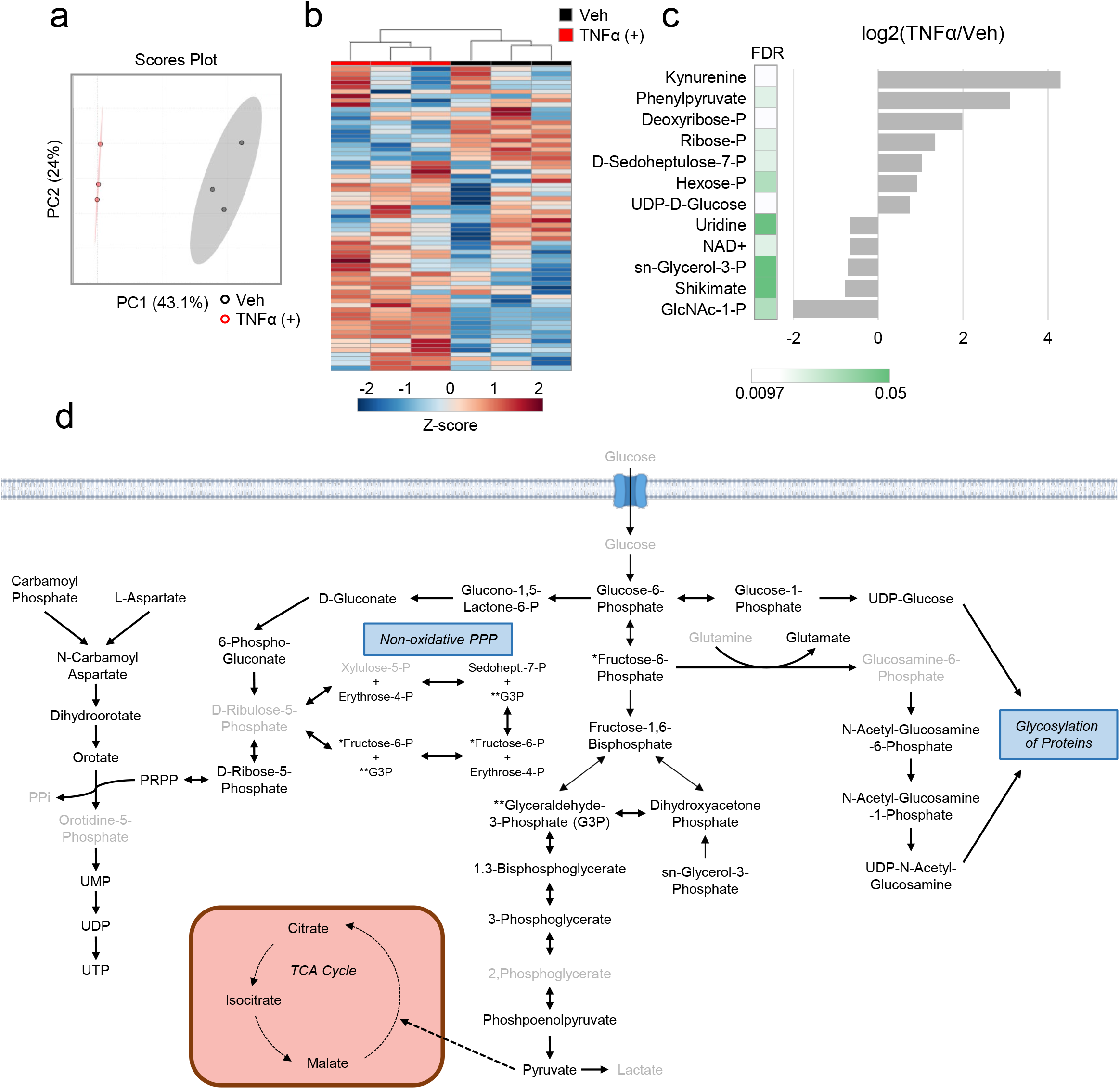
TNFα treatment induces broad metabolic changes. **(a-c)** HFFs were treated with TNFα (10 ng/mL, red) or vehicle (black) for 24 hr. Metabolites were extracted from cells and analyzed by LC-MS/MS as indicated in the materials and methods (n=3). **a** Principal component analysis (PCA) of metabolite data. **b** Hierarchical clustering of metabolite data depicted as Z-scores from min (blue) to max (red). **c** Significantly changed metabolites upon TNFα treatment. **d** Schematic representing metabolites in pathways of interest. Metabolites depicted in black are experimentally detected, whereas those in grey text are not detected. Solid lines represent a direct metabolic conversion while dashed lines represent an indirect conversion. Single headed arrows represent an irreversible reaction, double headed arrows represent a reversible reaction.

Given TNFα’s impact on glycolytic and pentose phosphate metabolite abundances, we more thoroughly investigated the impact of TNFα treatment on metabolites from these pathways (Fig. 1d) over a time course. TNFα treatment substantially increased several glycolytic metabolite pools over multiple time points (Fig. 2a, Supplementary Table 2). Four hours after TNFα treatment, the levels of fructose-1,6-bisphosphate, whose production is one of the rate-determining steps of glycolysis^26^, more than doubled. Other glycolytic pools were induced by TNFα treatment at multiple time points including fructose-6-phosphate, dihydroxyacetone phosphate, and glyceraldehyde-3-phosphate (Fig. 2a). Similarly, UDP-glucose of the hexosamine pathway was induced at every time point analyzed (Fig. 2a). For pentose phosphate metabolites: sedoheptulose-7-phosphate was only induced at 24 hours post-TNFα treatment, whereas erythrose-4-phosphate and ribose-phosphate were induced at all time points analyzed (Fig. 2a).

**Figure 2.**
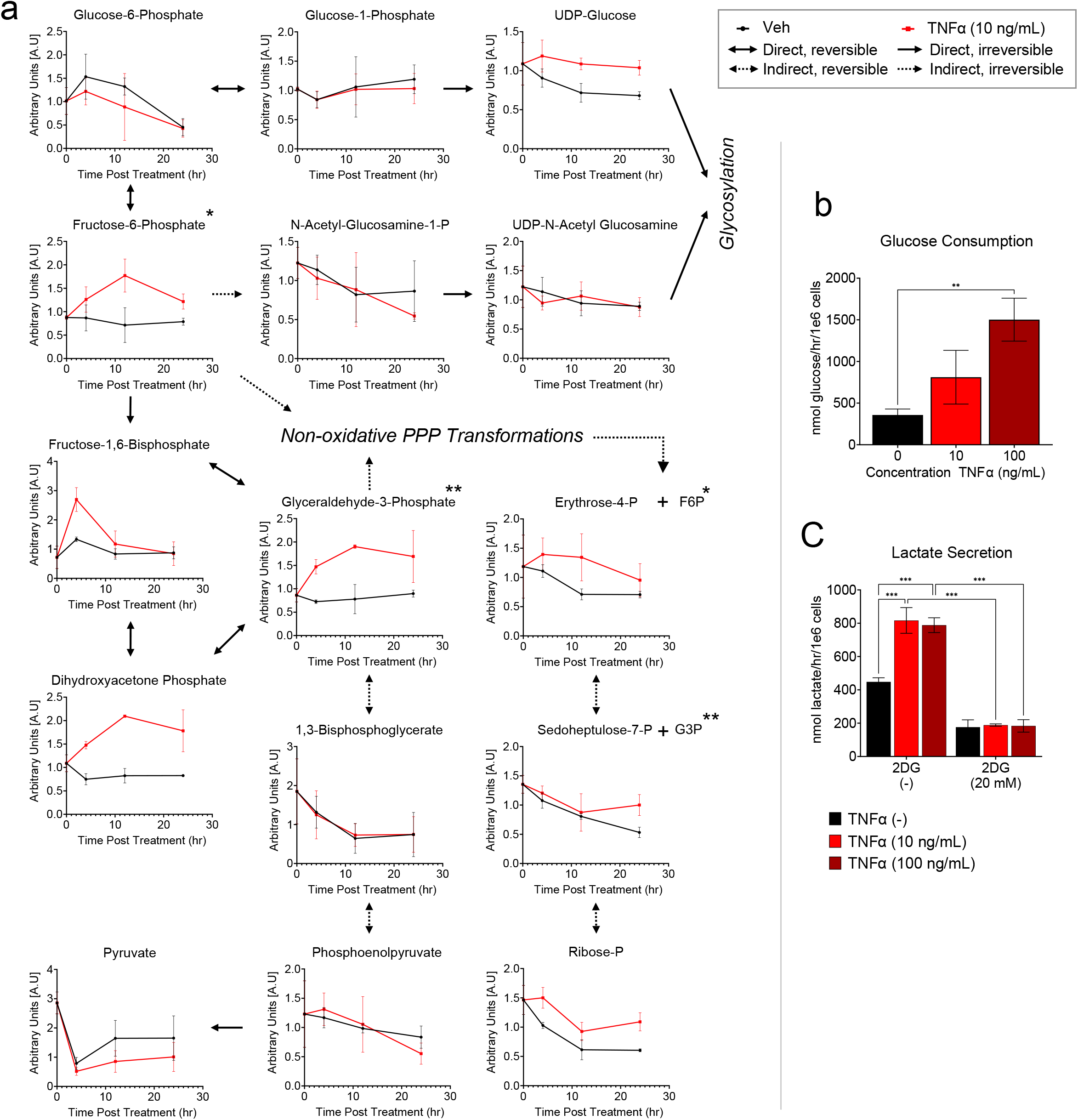
TNFα treatment induces glycolytic flux. **a** HFFs were treated with TNFα (10 ng/mL, red) or vehicle (black) for 0, 4, 12 and 24 hr prior to extraction and LC-MS/MS analysis (mean ± SD plotted, n=3). Solid arrows represent direct metabolite conversions while dashed arrows represent indirect conversions. Single-headed arrows represent irreversible conversions while double-headed arrows represent reversible conversions. **(b/c)** HFFs were treated with TNFα (10 ng/mL, light red or 100 ng/mL, dark red) or vehicle (black) for 24 hr. Media harvested from each sample were analyzed for **b** glucose consumption and **c** lactate secretion. (mean ± SD, n=3, with FDR adjusted ANOVA p-values, *p<0.05, **p<0.01, ***p<0.001).

The elevation of several glycolytic intermediates suggested that TNFα-treatment induced glycolysis. To test this possibility, we analyzed the impact of TNFα on glucose consumption and lactate secretion. Both glucose consumption and lactate secretion were substantially induced by TNFα treatment (Fig. 2b & c). Basal lactate secretion and TNFα-induced lactate secretion were severely attenuated by treatment with the glycolytic inhibitor, 2-deoxyglucose (2DG) (Fig. 2c). Collectively, our data indicate that TNFα treatment induces glycolytic activation as part of a broad metabolic remodeling.

### TNFα-mediated glycolytic activation is important for its anti-viral activity

To examine how glycolytic inhibition impacted TNFα-induced metabolic modulation, we analyzed the metabolomic impact of 2DG co-treatment with TNFα via LC-MS/MS (Supplementary Table 3). PCA indicated that the largest amount of data variance, i.e., 37.1% of the variance associated with PC1, separated all 2DG treated from the non-2DG treated samples regardless of TNFα treatment (Fig. 3a). In the non-2DG treated samples, TNFα treated samples were separated from non-TNFα treated samples along PC2 (Fig. 3a). In contrast, co-treatment with 2DG largely collapsed this separation between TNFα treated and non-TNFα treated samples along PC2 (Fig. 3a). Similarly, hierarchical clustering of the metabolic data primarily separated samples based on whether they were 2DG treated or not (Fig. 3b). Further, as would be expected, 2DG treatment reversed the TNFα-mediated induction in glycolytic pools sizes (Fig. 3c, Cluster I, & 3c) and lactate secretion (Fig. 2c).

**Figure 3.**
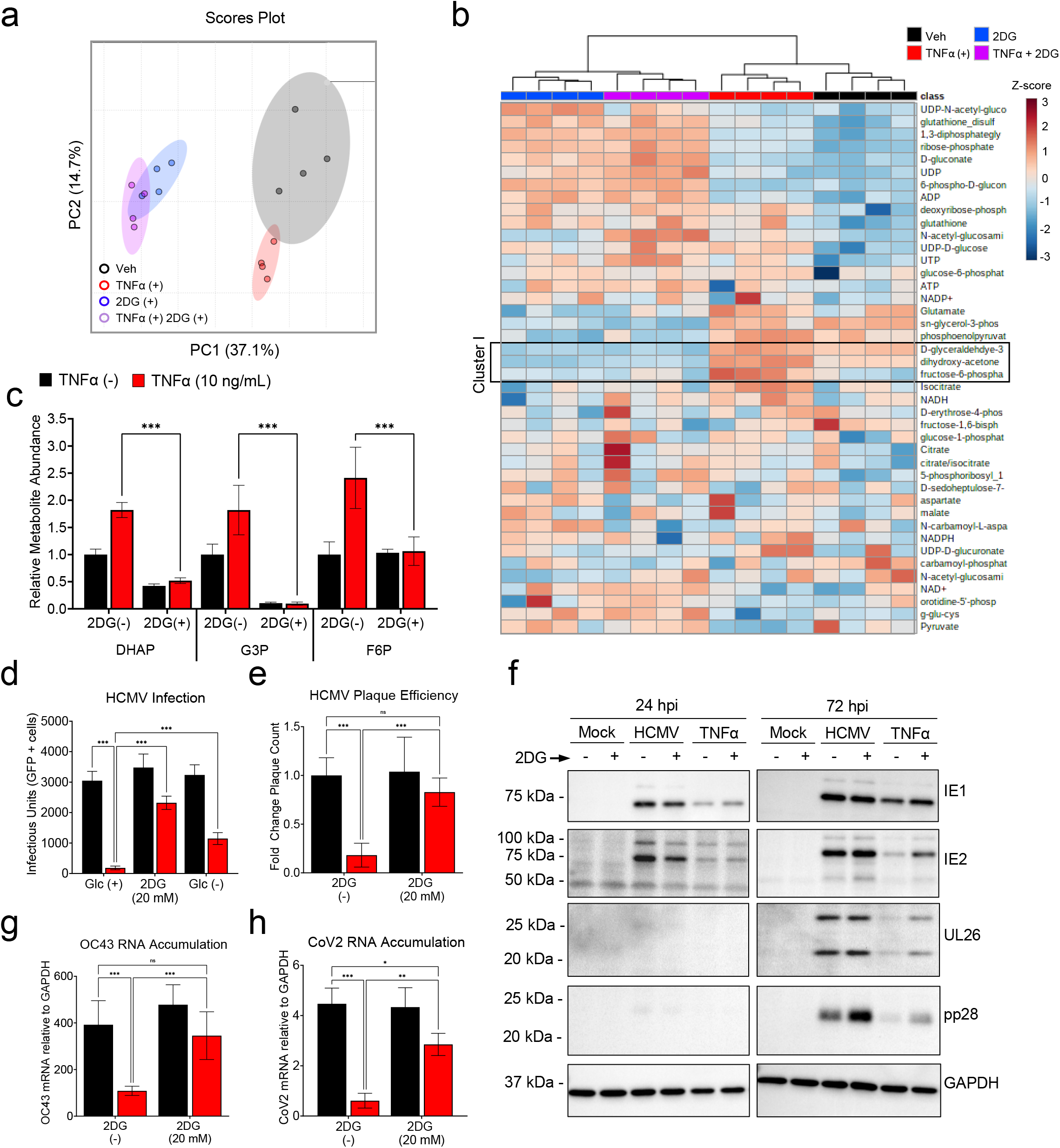
Glycolytic flux is required for TNFα to induce its anti-viral state. **(a-c)** HFFs treated with TNFα (10 ng/mL, red), 2DG (20 mM, blue), TNFα+2DG (purple), or vehicle (black) for 24 hr (n=4). Metabolites were extracted from cells and analyzed by LC-MS/MS (mean ± SD, n=4). Metabolite data analyzed via **a** PCA or **b** Hierarchical clustering with metabolite values depicted as Z-scores from min (blue) to max (red). **c** Relative metabolite abundance. **d** HFFs treated with TNFα (10 ng/mL, red) in the presence or absence of glucose (Glc ±) or 2-deoxyglucose (2DG, 20 mM) for 24 hr prior to a media change with fresh viral adsorption media containing glucose, but without inhibitors, and HCMV-GFP (MOI=0.5). Infected cells were identified by GFP expression 24 hr post-infection and quantified (n=12, mean ± SD). **e** HFFs pretreated as indicated for 24 hr prior to media change with fresh viral adsorption media containing glucose, but without inhibitors, and a known number of HCMV-GFP infectious particles. Viral plaques were quantified (n=6, relative mean ± SD). **f** Cells were pre-treated as in (**e**), infected with HCMV (MOI=3), and processed for western analysis at the indicated times post-infection. **(g-h)** Cells were pretreated as in (**e**), infected with **g** OC43 (MOI=3) or **h** SARS-CoV2 (MOI=0.01) and harvested 48 hr post-infection for RT-qPCR analysis (mean ± SD, n=6 (**g**) or n=3 (**h**)). p-values were calculated using two-way ANOVA and FDR corrected; *p<0.05, **p<0.01, ***p<0.001.

To determine how TNFα-mediated glycolytic activation contributes to its ability to induce the anti-viral state, we assessed how glycolytic inhibition impacted TNFα’s ability to limit viral infection. Pretreatment of cells with TNFα largely blocked the ability of HCMV to initiate infection (Fig. 3d & e). TNFα pretreatment in the presence of 2DG largely restored the ability of HCMV to infect cells (Fig. 3d & e). We also found that glucose starvation during TNFα pretreatment significantly increased the ability of HCMV to infect cells compared to TNFα pretreatment in glucose-containing medium (Fig. 3d). Pretreatment with TNFα prior to infection at a high multiplicity of infection (MOI=3.0) substantially reduced the accumulation of HCMV proteins throughout infection (Fig. 3f). Similar to the initiation experiments, TNFα pretreatment in the presence of 2DG partially restored HCMV protein expression (Fig. 3f). These results suggest that TNFα’s ability to promote an anti-viral cell state against HCMV relies on glucose availability and glycolytic flux.

Given that TNFα is broadly anti-viral, we next sought to determine if TNFα’s effects on glycolysis was specific to limiting HCMV infection, or if a similar phenotype could be observed in the context of evolutionary divergent viruses. To address this issue, we analyzed the replication of two β-Coronaviruses, OC43 and SARS-CoV-2, in cells that had been pretreated with TNFα in the presence or absence of 2DG. TNFα pretreatment was sufficient to restrict the RNA accumulation of both OC43 and SARS-CoV-2 (Fig. 3g & h). TNFα pretreatment in the presence of 2DG largely restored OC43 and SARS-CoV-2 RNA accumulation (Fig. 3g & h). These data align with the results from HCMV infection and support that TNFα treatment broadly requires glycolysis for its anti-viral activity.

### Hexokinase 2 contributes to TNFα-mediated glycolytic activation

To obtain a complimentary picture of TNFα-induced metabolic changes and how glycolysis affects these changes, we analyzed the impact of TNFα treatment in the presence or absence of 2DG on the proteome. The abundances of 3,780 unique proteins were identified in this study (Supplementary Table 4). From this list, we extracted those involved in metabolism to study the differences in the abundance of metabolic enzymes between vehicle and TNFα treated cells. Of the 562 metabolic enzymes detected in our study, 40 were significantly more abundant as a result of TNFα treatment (Supplementary Table 4.1). Hierarchical clustering (Fig. 4a) and PCA (Supplementary Fig. 1) of this subset of metabolic enzymes showed that TNFα-treated samples segregate from vehicle-treated samples suggesting that TNFα treatment substantially impacts the expression of metabolically categorized proteins. Major contributors to this shift included SLC2A1 (the ubiquitous GLUT1 glucose transporter), and hexokinase-2 (HK2), the muscle predominant form of hexokinase (Fig. 4a & b), both of which are proteins involved in glucose metabolism/glycolysis. GLUT1 was induced ∼5-fold, and HK2 was induced ∼2.5 fold in TNFα-treated samples relative to vehicle (Fig. 4c). In agreement with our proteomics data, HK2 mRNA was induced upon TNFα treatment (Fig. 4d), as was GLUT1 mRNA (Fig. 4e). Immunoblot analysis showed HK2 and GLUT1 proteins were ∼40% and ∼90% more abundant in TNFα treated cells, respectively (Fig. 4g). No other glycolytic enzymes were significantly increased as a result of TNFα treatment, however in contrast PFK-M and TIGAR levels were significantly reduced by TNFα treatment (Fig. 4b & c). We also sought to examine if TNFα could be inducing HIF1α expression given its role as a glycolysis activator^27^. While Hif1α was not detected in the proteomics data set after 24 hours of TNFα treatment, its mRNA levels were induced 2-fold at 8 hours post-TNFα treatment relative to vehicle-treated samples (Fig. 4f).

**Figure 4.**
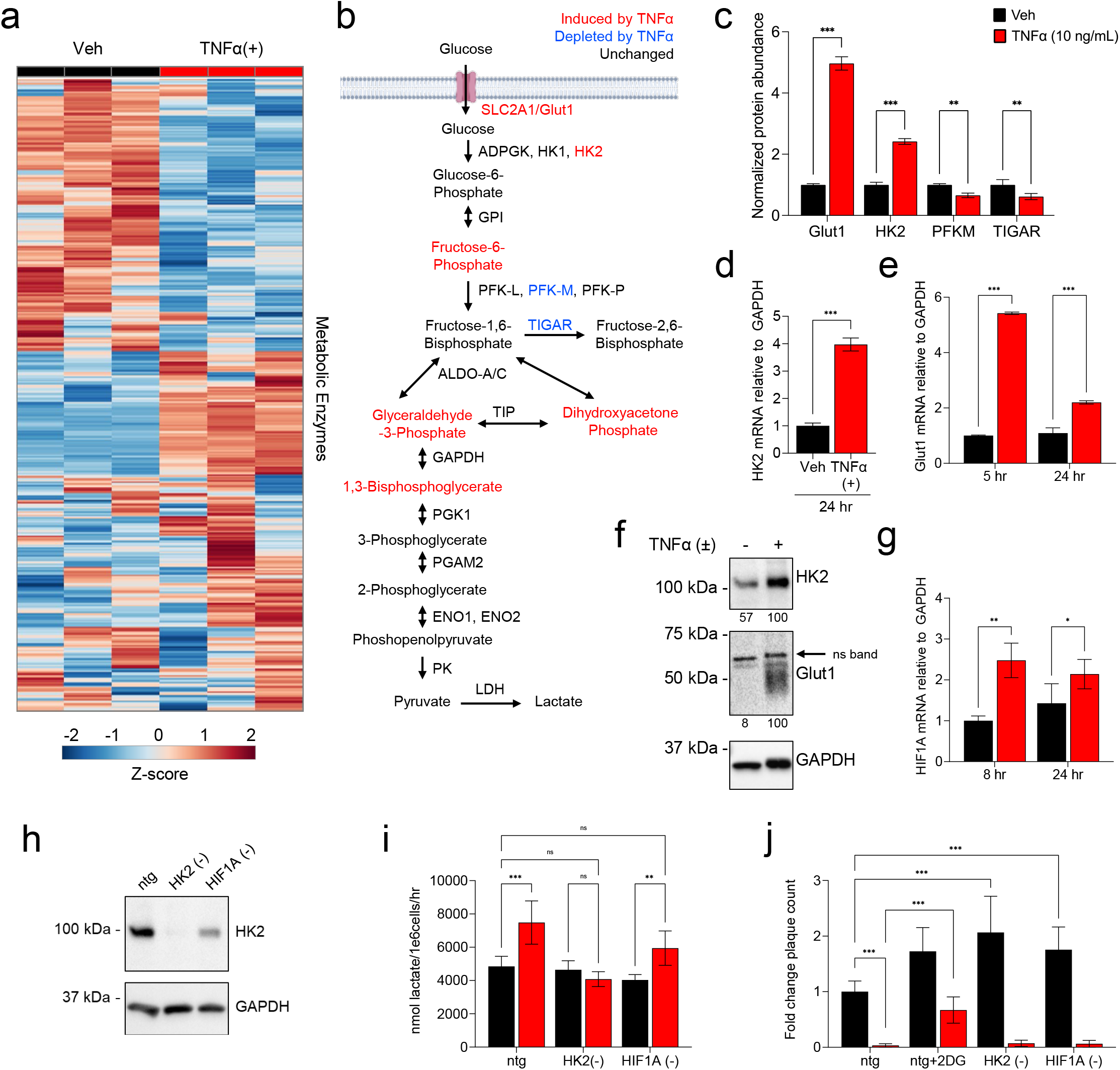
TNFα treatment induces HK2 expression to support glycolytic flux. **(a-f)** HFFs were treated with TNFα (10 ng/mL, red) or vehicle (black) for 24 hr, unless otherwise indicated. **(a-c)** Cells were harvested for proteomics analysis. **a** Hierarchical clustering of proteins involved in metabolism. Protein abundance values are depicted as Z-scores from min (blue) to max (red). **b** Schematic of metabolites and enzymes involved in glycolysis significantly dysregulated by TNFα treatment. Red = significantly more abundant, blue = significantly less abundant. **c** Proteomics data for Glut1, HK2, PFK-M and TIGAR normalized to vehicle-treated (mean ± SD, n=3). (**d/e/g/f)** Cells were harvested at the indicated times post TNFα treatment for RT-qPCR analysis of **d** HK2, **e** Glut1 and **g** Hif1α mRNA or **f** western analysis as indicated. Band intensities quantified and normalized to the intensity of their respective GAPDH bands. **(h-j)** Cells treated with CRISPR Cas9-RNP containing guides for HK2, Hif1a, or a non-target guide (ntg) to generate knockout (KO) cell lines. KO cells harvested for **h** western analysis as indicated. KO cells treated with TNFα (10 ng/mL, red) or vehicle (black) for 24 hr, **i** media analyzed for lactate secretion. **j** KO cells pre-treated with 2DG (20 mM, black) and TNFα (10 ng/mL) as indicated prior to media change with fresh viral adsorption media, without inhibitors, containing a known number of HCMV-GFP infectious particles. Viral plaques were quantified and plotted as indicated. **d** mean ± SD, n=3, two-tailed unpaired ttest, *p<0.05, **p<0.01, ***p<0.001. **c/e/g** mean ± SD, n=3; **i** mean ± SD, n=4; **j** mean ± SD n=15 with FDR-adjusted p-values determined using 2-way ANOVA followed by two-stage step-up method of Benjamini, Krieger and Yekutieli *p<0.05, **p<0.01, ***p<0.001.

Given that HK2 is the muscle-specific hexokinase isoform, and is not thought to be substantially expressed in fibroblasts, we sought to determine the importance of TNFα-induced expression of this isoform for glycolytic activation. We targeted HK2 and HIF1α via CRISPR-Cas9 to assess their contributions to TNFα-mediated glycolytic activation and induction of an anti-viral cell state. RNP-based Cas9 delivery successfully inactivated HK2, resulting in the loss of HK2 protein accumulation (Fig. 4h). While Hif1α protein could not be detected by western blot, genomic ablation of the HIF1A gene was confirmed via genomic sequence analysis, with a knockout-score of 89% (Supplementary Fig. 2). Notably in HIF1A targeted cells, HK2 protein expression was somewhat reduced, raising the possibility that HK2 could be regulated by Hif1α (Fig. 4h). We first tested the effect of HK2 and HIF1 knockout (KO) on lactate secretion compared to non-targeting guide (ntg) control cells. HK2 KO cells treated with TNFα did not exhibit an induction of lactate secretion relative to vehicle treated HK2 KO cells (Fig. 4i), suggesting that HK2 accumulation is important for TNFα-mediated glycolytic activation. In contrast, lactate secretion in HIF1A KO cells treated with TNFα was modestly induced compared to vehicle treated HIF1A KO cells (Fig. 4h), suggesting that Hif1α may largely be dispensable for TNFα-mediated glycolytic activation.

We next tested if HK2 or Hif1α are necessary for TNFα to promote an anti-viral state. As expected, control ntg cells pretreated with TNFα in the presence of the glycolytic inhibitor 2DG are more permissive to HCMV infection (Fig. 4j). Interestingly, non-TNFα treated HK2 KO and HIF1A KO cells showed an increase in HCMV plaque formation relative to the vehicle pretreated control ntg cells, suggesting that HIF1A and HK2 might restrict the initiation of HCMV infection in the absence of TNFα (Fig. 4j). However, HK2 KO and HIF1A KO cells pretreated with TNFα fully restricted HCMV initiation of infection (Fig. 4j), indicating these factors are not necessary for TNFα’s anti-viral activity, and thereby suggesting that enough glycolysis occurs in their absence to enable TNFα-mediated viral inhibition. Our data suggest that while HK2 and Hif1α are not essential for TNFα’s anti-viral activity, they may play a role in restricting HCMV infection that is independent of TNFα signaling.

### Glycolytic inhibition attenuates the accumulation of intrinsic anti-viral proteins upon TNFα treatment

To get a more global picture of how glycolytic inhibition affects the TNFα-induced anti-viral state, we reexamined the proteomic data set to compare the impact of TNFα treatment in the presence and absence of 2DG (Supplementary Table 4). PCA of the total proteomics data set show that TNFα-treated samples in the presence or absence of 2DG segregated from non-TNFα treated samples largely along PC2, whereas 2DG-treated samples segregated from non-2DG treated samples along PC1 (Fig. 5a). Notably, co-treatment with 2DG and TNFα blunted the TNFα-induced response as indicated by a much-reduced shift along PC2. In contrast, co-treatment with 2DG and TNFα induced a larger leftward shift along PC1 than was observed with 2DG treatment alone (Fig. 5a). These data suggest that 2DG treatment blunts the impact of TNFα treatment on the proteome, and further that TNFα and 2DG co-treatment accelerates the proteomic impact over 2DG alone.

**Figure 5.**
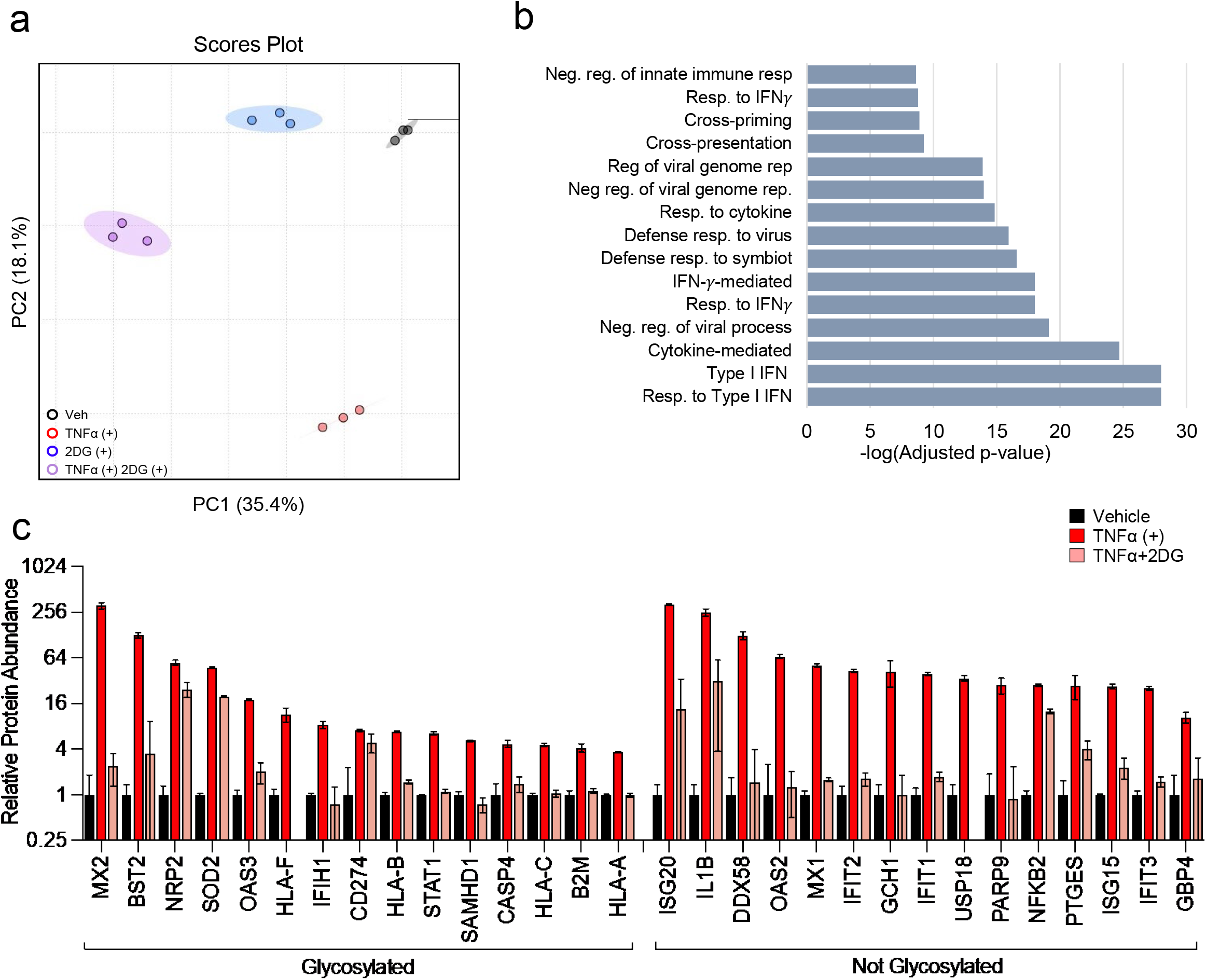
TNFα requires glycolysis to promote expression of intrinsic anti-viral factors and immunoregulatory proteins. **(a-c)** HFFs were treated with TNFα (10 ng/mL, red), 2DG (20 mM, blue), TNFα+2DG (purple) or vehicle (black) for 24 hr. Cells were harvested and analyzed by LC-MS/MS for protein abundance. **a** PCA of relative protein abundance from proteomic results. **b** Ontology analysis of TNFα-induced proteins that were significantly decreased upon co-treatment with 2DG. **c** Abundance of proteins belonging to the Response to Cytokine ontology [GO:0034097] (mean ± SD, n=3).

To further explore the impact of 2DG treatment on TNFα-induced proteomic changes, we analyzed the proteins whose abundances were significantly depleted or induced by co-treatment with 2DG and TNFα relative to TNFα alone (Supplementary Table 4.2 and 4.3, respectively). Gene Ontology (GO) analysis indicated that proteins significantly up-regulated in response to 2DG+TNFα co-treatment relative to TNFα treatment alone belonged to a diverse array of pathways including receptor-mediated endocytosis, the unfolded protein response, and various oxidative and metabolic stress responses (Supplementary Table 4.3, Supplementary Figure 3). In contrast, the pathways associated with proteins that exhibited substantially lower levels during 2DG and TNFα co-treatment relative to TNFα alone shared a common theme, they are important to the innate immune antiviral response (Fig. 5b, Supplementary Table 4.2). These ontological pathways included responses to IFN, defense responses to viruses, and regulation of viral genome replication (Fig. 5b). Approximately 33% of these proteins reduced in 2DG and TNFα co-treated cells included genes within the ontological pathway ‘response to cytokine signaling’, e.g., IFIs, ISGs, as well as other critical viral defense genes, e.g., MX1, OAS53, SAMHD1, and global innate regulators, e.g., STAT1 (Fig. 5c, Supplementary Fig. 4). Additionally, 43% of these substantially down-regulated proteins are glycosylated (Fig. 5c, Supplementary Fig. 4). Collectively, these data indicate that restricting glycolysis during TNFα treatment prevents the accumulation of proteins involved in the anti-viral response, a substantial proportion of which are glycosylated.

### TNFα treatment induces the accumulation of glycosyl precursors and glycosylation enzymes

Glucose provides the subunits necessary for protein glycosylation, which is critical for the stability of numerous glycoproteins. Given the observation that TNFα activates glycolysis, and that inhibition of glycolysis results in decreased accumulation of several glycosylated anti-viral proteins (Fig. 5c), we sought to determine the extent to which TNFα impacts the generation of glycosyl precursors. Our proteomics data show three enzymes involved in the production and utilization of glycosyl precursors UDP-glucose (UDP-Glc) and UDP-N-Acetyl Glucosamine (UDP-GlcNAc) were more abundant in TNFα-treated cells (Fig. 6a). These enzymes include UTP-glucose-1-phosphate uridylyl transferase (UGP2), which catalyzes the conversion of glucose-1-phosphate into UDP-Glc^28^; dolichyl-phosphate beta-glucosyltransferase (ALG5), which initiates biosynthesis of lipid-linked oligosaccharides for glycosylation in the ER membrane using UDP-Glc^29^; and alpha-1,3-mannosyl-glycoprotein 2-beta-N-acetylglucosamine transferase (MGAT1), which promotes biosynthesis of N-Glycans for glycosylation using UDP-GlcNAc as a substrate^30^ (Fig. 6a). We find that UDP-Glc is ∼3-fold more abundant in cells treated with TNFα, but UDP-GlcNAc pools are unchanged (Fig. 6b). To examine the turnover kinetics of these pools, we labeled vehicle or TNFα treated cells with U-^13^C-glucose over time. TNFα treatment induced the accumulation rate of ^13^C-UDP-Glc and ^13^C -UDP-GlcNAc isotopologues, while increasing the rate of ^12^C isotopologue disappearance (Fig. 6c & d), consistent with activation of these pathways. These data suggest that TNFα treatment induces activation of UDP-sugar metabolism to support protein glycosylation.

**Figure 6.**
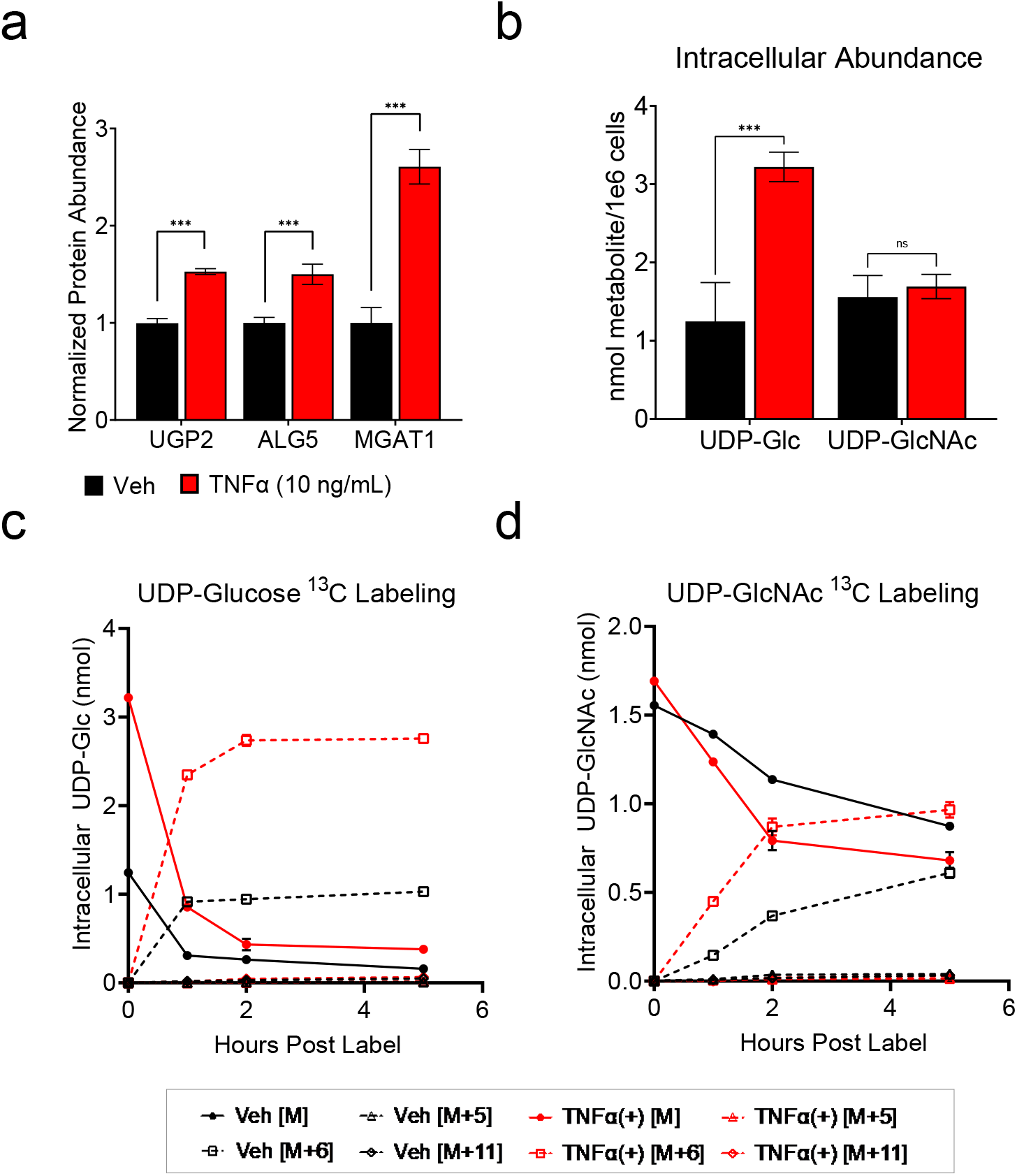
TNFα treatment induces glycosylation precursor turnover. HFFs were treated with TNFα (10 ng/mL, red) or vehicle (black) for 24 hr. **a** Cells harvested and analyzed by LC-MS/MS for protein abundance. Normalized protein abundance of UTP-glucose-1-phosphate uridylyl transferase (UGP2), Dolichyl-phosphate beta-glucosyltransferase (ALG5), and Alpha-1,3-mannosyl-glycoprotein 2-beta-N-acetylglucosamine transferase (MGAT1) (mean ± SD, n=3). **b** Metabolites extracted from cells and analyzed by LC-MS/MS for UDP-Glucose (UDP-Glc) and UDP-N-Acetyl Glucosamine (UDP-GlcNAc) intracellular concentrations (mean ± SD, n=3). **(c/d)** HFFs treated with TNFα (10 ng/mL, red) or vehicle (black) for 19 hours in media containing unlabeled ^12^C-glucose prior to media change with media containing U-^13^C-glucose. Cellular extracts were harvested at t=0, 1, 2 and 5 hr post-label addition. **C** UDP-Glc or **d** UDP-GlcNAc intracellular isotopologue abundances were quantified by LC-MS/MS (mean ± SD, n=3). Solid lines represent intracellular abundances of unlabeled ^12^C metabolite species, dashed lines represent intracellular abundances of ^13^C-labeled metabolite species. M refers to the ^12^C unlabeled species and M+n represents detection of a ^13^C-labeled species where n represents the number of additional mass units detected by mass spec **a/b** FDR-adjusted p-values were determined using 2-way ANOVA followed by two-stage step-up method of Benjamini, Krieger and Yekutieli; ns = not significant, *p<0.05, **p<0.01, ***p<0.001.

### Glycosylation is important for the TNFα-induced anti-viral state

To dissect how various metabolic pathways contribute to TNFα’s ability to induce the anti-viral cell state, we pretreated cells with a variety of inhibitors that attenuate metabolic pathways adjacent to glycolysis in the presence or absence of TNFα (Fig. 7a). Two inhibitors of pyrimidine biosynthesis [*N*-phosphonacetyl-L-aspartate (PALA)^31^ and vidoflumidus (Vid)^32^], and two pentose phosphate pathway inhibitors [6-aminonicotinamide (6-AN)^33^ and N3-pyridyl thiamine (N3-PT)^34^] failed to rescue viral replication when co-pretreated with TNFα (Fig. 7a &7b). In contrast, co-pretreatment of TNFα with Benzyl-α-GalNAc (BGNAc), an inhibitor of *O*-linked glycosylation^35, 36^ largely rescued the ability of HCMV to initiate infection. The magnitude of BGNAc rescue was indistinguishable from the rescue observed with 2DG co-pretreatment (Fig. 7b). These results support a model in which protein glycosylation plays a critical role in TNFα’s ability to induce the anti-viral cell state.

**Figure 7.**
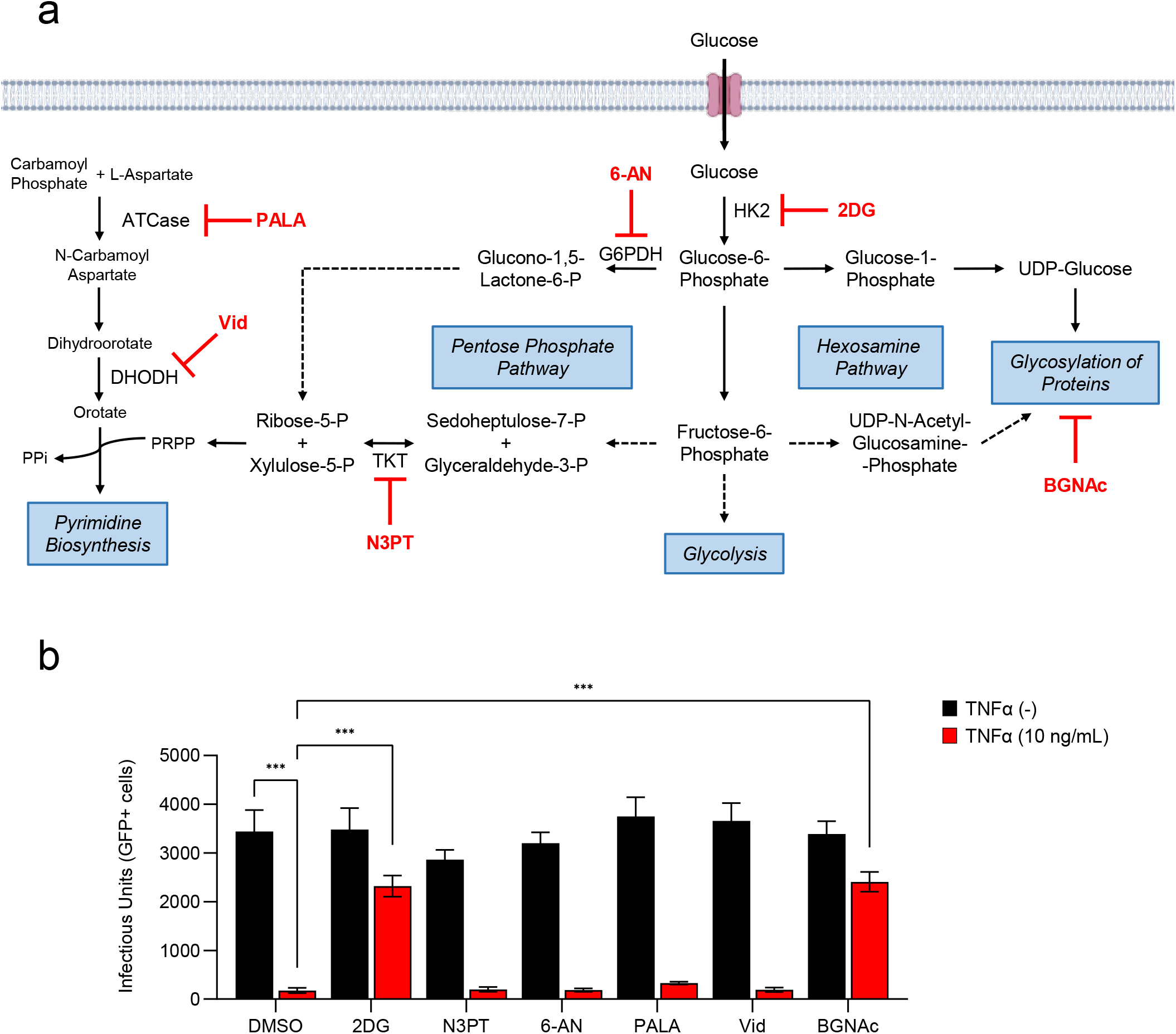
Inhibition of glycolysis and glycosylation restricts TNFα’s ability to induce the anti-viral state. **a** Schematic of glucose metabolism and inhibitors tested (red). **b** HFFs treated with TNFα (10 ng/mL, red) or vehicle (black), in the presence or absence of 2-Deoxyglucose (2DG, 20 mM), N3-pyridyl thiamine (N3PT, 50 nM), 6-aminonicotinamide (6-AN, 200 nM), *N*-phosphonacetyl-L-aspartate (PALA, 150 uM), vidiflumidus (vid, 300 nM) or Benzyl-α-GalNAc (BGNAc,15 mM) or DMSO for 24 hr prior to a media change with fresh viral adsorption media containing glucose, but without inhibitors, and HCMV-GFP (MOI=0.5). Infected cells were quantified by GFP expression measured at 24 hr post-infection (n ≥6, mean ± SD). p-values were calculated using two-way ANOVA with Tukey’s multiple comparison test; *p<0.05, **p<0.01, ***p<0.001.

## Discussion

TNFα pretreatment promotes an anti-viral cell state that prevents replication of diverse viral families^8-10^. However, many questions remain about the functional requirements and cellular activities responsible for limiting viral infection. In this study, we found that TNFα treatment induces broad metabolic changes, including activating glycolysis and inducing UDP-sugar biosynthesis. Further, glycolysis was necessary for the accumulation of several glycosylated anti-viral proteins. TNFα treatment also increased the accumulation of glycosyltransferases such as ALG5 and MGAT1. Inhibiting glycolysis or glycosylation during TNFα pretreatment resulted in the loss of TNFα’s ability to attenuate viral replication. Together, these data suggest a mechanism wherein TNFα activates glycolysis and glycosylation as essential components of instituting an anti-viral state.

A key aspect of TNFα-induced glycolytic activation was the induction of HK2 expression, which was subsequently found to be important for TNFα-mediated glycolytic activation (Fig. 4). Previously, TNFα has been reported to induce HK2 in a murine skeletal muscle cell line^37^, but little is known about how HK2 contributes to TNFα-associated activities. HK2 is canonically thought of as the muscle-specific hexokinase isoform, although it’s expression has also been found to be induced in various tumor types^38^. In tumors it is thought to be important for cancer-cell survival through modulation of metabolic-associated stress responses^39^. While our results indicate that TNFα induces HK2 expression, which we found to be important for TNFα-induced glycolytic activation, its deletion did not restore viral infection in the face of TNFα treatment (Fig. 4j). This suggests that HK2 is not essential for TNFα-mediated inhibition of HCMV plaque formation, and that the amount of glycolysis that occurs in its absence is sufficient to support TNFα’s antiviral activity. However, it was notable that inactivation of HK2 significantly increased HCMV-mediated plaque formation in the absence of TNFα treatment (Fig 4j), suggesting that HK2 could be playing a role in intrinsic anti-viral defense. Future work should explore this possibility, as well as the possibility that HK2 induction is contributing to other aspects of TNFα-modulated innate immunity, for example, processing or presentation of viral antigens.

Hif1α is a central regulator of glycolytic gene expression and was implicated in TNFα-mediated induction of HK2 in murine skeletal muscle cells^37^, raising the possibility that Hif1α is important for TNFα-mediated induction of glycolysis. Our data suggest that Hif1α is dispensable for TNFα-mediated induction of lactate secretion and inhibition of HCMV replication (Fig 4i & 4j). Separately, our data show that cells lacking Hif1α are more permissive to infection, suggesting Hif1α may play a role in limiting the initiation of infection. Consistent with this finding, Hif1α was recently shown to attenuate HCMV replication in human fibroblasts^40^. The anti-viral phenotype associated with Hif1α-mediated metabolic regulation is intriguing, yet the mechanisms responsible for TNFα-induced glycolysis still require elucidation.

Our data show that TNFα induces changes to several metabolic pools. Our efforts focused primarily on glycolysis, as a number of metabolites upregulated by TNFα were involved in glucose metabolism. However, the metabolite most strongly induced by TNFα treatment was kynurenine (Fig. 1c). Kynurenine is a tryptophan-related metabolite that can also be considered part of the NAD^+^ biosynthetic pathway. Kynurenine accumulates in a variety of inflammatory conditions and is induced during Human Immunodeficiency Virus (HIV) infection^41, 42^. Others have described pro-viral effects of kynurenine on HCMV replication^40^, suggesting a potential complex relationship with infection that requires further analysis with respect to its roles in inflammation, intrinsic immunity and during viral infection.

In addition to activation of glycolysis, we find that TNFα induces the accumulation UDP-Glucose (UDP-Glc) and stimulates the glucose-mediated labeling of both UDP-Glc and UDP-N-Acetyl-Glucosamine (UDP-GlcNAc) (Fig. 6). These nucleotide sugars are molecular substrates for glycosylation reactions that are critical for a variety of cellular activities including the stability of many proteins^43, 44^, as well as immune cell differentiation and activation^45, 46^. Inhibition of O-linked glycosylation phenocopied glycolytic inhibition in that it prevented TNFα-mediated inhibition of HCMV infection, highlighting the importance of glycosylation to TNFα’s anti-viral activity (Fig 7). A number of glycosylated anti-viral effector proteins failed to accumulate upon TNFα treatment in the face glycolytic inhibition, including MX2, BST2, OSA3 and STAT1, as did various glycosylated components of the adaptive immune response including B2M and HLA-A/B/C/F (Fig. 5). These data are consistent with a model in which TNFα treatment drives UDP-sugar production to support the glycosylation and stable expression of anti-viral effector proteins.

That the expression of so many diverse anti-viral effector proteins was impacted by glycolytic inhibition in the face of TNFα treatment likely explains our findings that glycolysis is necessary to attenuate both HCMV and coronavirus replication. HCMV and the coronaviruses tested, OC43 and SARS-CoV2, possess extremely different viral life cycles, e.g., nuclear DNA replication versus cytoplasmic RNA replication, but the anti-viral effectors downregulated by glycolytic inhibition target multiple aspects of various viral infections. These intrinsic anti-viral proteins included MX2, which can block nuclear capsid transport^47^; OAS3, whose activity can result in viral RNA degradation^48^; and BST2, which can tether viruses to membranes for degradation^49, 50^. Collectively, our data indicate that TNFα-mediated metabolic remodeling is broadly important for its anti-viral activity.

A number of non-glycosylated anti-viral proteins failed to accumulate upon TNFα treatment when glycolysis was inhibited (Fig 5), which could reflect a secondary dependence on the accumulation of a crucial glycosylated protein. STAT1, for example, is glycosylated and fails to accumulate upon TNFα treatment in the presence of glycolytic inhibition (Fig 5). Given STAT1’s importance in the transcription of numerous anti-viral effectors^51^, it would be predicted that its loss would deplete the expression of glycosylated and non-glycosylated anti-viral effector proteins. Potential secondary effects of losing STAT1 notwithstanding, it is still plausible that in addition to providing UDP-sugar subunits to support glycosylation, glycolysis contributes to TNFα’s anti-viral activity via other downstream metabolic activities. In this regard, many questions still remain about the molecular fate of glycolytically-derived carbon and how these pathways could contribute to TNFα’s anti-viral activity.

Our results indicate that TNFα-induced metabolic remodeling is important for its ability to promote an anti-viral state. However, many questions remain about other potential metabolic requirements for cytokine-induced intrinsic viral defense. What other cytokine-driven metabolic activities are important for limiting viral infection? What other aspects of innate immunity require specific metabolic activities, e.g., antigen processing and presentation? Is there a common anti-viral metabolic or glycosylation program induced by diverse anti-viral cytokines, e.g., TNFα, IFNγ, IFNα, etc. Also, given that viruses usurp cellular metabolic resources for their replication, are there cytokine induced anti-viral mechanisms that prevent pathogens from accessing cellular metabolic resources? The answers to these questions will likely shape our understanding of an important host pathogen interaction, that is, the metabolic regulation associated with intrinsic immunity in the face of viral infection and the associated contributions to preventing viral pathogenesis.

## Materials and Methods

### Cell Culture, Viruses and Viral infection

Telomerase-transduced Human Foreskin Fibroblasts (HFFs) and MRC5 lung fibroblasts were cultured in Dulbecco’s modified Eagle serum (DMEM; Invitrogen #11965118) supplemented with 10% (v/v) fetal bovine serum (FBS), and 1% penicillin-streptomycin (Pen-Strep; Life Technologies #15140-122) at 37≥C in a 5% (v/v) CO2 atmosphere. VEROe6 cells (ATCC, #30-2003) were cultured in Minimum Essential Medium Eagle serum (MEM, Invitrogen) supplemented with 10% (v/v) fetal bovine serum (FBS), 1X GlutaMAX (Invitrogen #35050-061), and 1% penicillin-streptomycin at 37≥C in a 5% (v/v) CO2 atmosphere. HFFs were transduced with ACE2 (HFF-ACE2) with lentivirus as previously described^52^.

The WT strain of HCMV used in experiments was BADwt, a Bacterial Artificial Chromosome (BAC) clone of AD169. GFP expressing AD169, referred to as WT-GFP, was BADsubUL21.5^53^. OC43 and SARS-CoV-2, Isolate Hong Kong/VM20001061/2020 (BEI Resources NR-52282) were cultured as previously described^52^. All experiments involving live SARSCoV-2 were conducted in a biosafety level 3 facility at the University of Rochester. Viral stocks of HCMV, OC43 and SARS-CoV2 were propagated and titered as described previously^52^ using a modified Reed & Muench TCID50 calculator from the Lindenbach lab^54^.

### Reagents and Preparation of Treatments

For all experiments where cells were treated with TNFα and/or inhibitors, cells were grown to confluence. Twenty four hours prior to infection, the medium was replaced with serum-free DMEM supplemented with 1% PenStrep and appropriate compounds as indicated.

TNFα (Human) was purchased from GoldBio (#1130-01-100) and suspended in sterilized water to 1 mg/mL concentration. Aliquots of 20 μL were stored at -80C. For experiments, TNFα was prepared in master mix solutions to 10 ng/mL concentration. 2-Deoxy-D-Glucose (2DG) was purchased from Millipore-Sigma (#D8375-5G) and prepared to 20 mM for experiments. 6-Aminonicotinamide (MedChem Express, Cat# HY-W01034) was suspended in DMSO to 10 mM and prepared to the indicated concentrations in culture medium. N3-pyridyl thiamine (MedChem Express, Cat# HY-16339, 5 mg) was suspended in DMSO to 8.5 mM and prepared to the indicated concentrations in culture medium. Sparfosic Acid (MedChem Express, Cat# HY-112732B) suspended in H2O to 10 mM and prepared to the indicated concentrations in culture medium. Vidofludimus (Cayman Chemicals, Item#18377 CAS# 717824-30-1) suspended in DMSO to 10 mM and prepared to the indicated concentrations in culture medium. Benzyl-2-acetamido-2-deoxy-α-D-galactopyranoside (Benzyl-alpha-GalNAc), (MedChem Express Cat# HY-129389) suspended in H2O to 25 mM using ultra sonication and prepared to indicated concentration in culture medium. All treatment medium were prepared in serum-free DMEM supplemented with 1% PenStrep unless otherwise indicated. Vehicle or control treatment was serum-free DMEM supplemented with 1% PenStrep.

### Plaque Efficiency Assay

For plaque efficiency assay, cells were grown to confluence in 12 well TC-treated Greiner plates (#82050-930) and treated as described above. Treatment medium was removed and the cell monolayer was infected with a low, known number of plaque forming units (PFUs) of AD169-GFP in 500 uL medium overnight. Following this adsorption period, medium was removed and replaced with 2 mL of a standard agarose gel overlay prepared with 2X concentrated DMEM (Invitrogen #12100046) supplemented with 10% FBS and 1% NuSieve Agarose (Lonza, 12001-722). Plaques were allowed to develop for 10 days and total number of plaques in each well was counted using a fluorescent microscope. For each indicated experimental condition, total plaque count of each well was normalized to the average plaque count of the control condition.

### Quantification of HCMV Infectious Units (GFP + cells)

HFFs were grown to confluence in 96-well TC-treated plates then treated for 24 hours in 100 μL indicated treatment medium. Treatment was removed and replaced with GFP-expressing HCMV viral inoculum (MOI = 0.5, 100 μL) in serum-free DMEM, high glucose, no glutamine, no phenol red medium (Invitrogen #31053036) supplemented with 1x Glutamax and 1% Pen-Strep for 24 hours. Medium was removed and replaced with PBS containing 1:1000 diluted Hoescht fluorescent stain (Thermo Fisher #33342). HFFs were then imaged using a Cytation 5 imaging reader (BioTek). Each well was imaged using a 4X magnification objective lens and predefined DAPI channel with an excitation wavelength of 377 nm and emission wavelength of 447 nm for nuclei count or GFP channel with an excitation wavelength of 469 nm and emission wavelength of 525 nm for GFP-expressing cells. Gen5 software (BioTek) was used to determine cell number by gating for objects with a minimum intensity of 3000, a size greater than 5 µm and smaller than 100 µm.

### Western Blot Analysis

Protein detection was performed as previously described^55^. Briefly, samples were harvested in SDS-lysis buffer (20 mM TrisHCl (VWR # 97061-794) pH7, 5% β-Mercaptoethanol (gibco #21985-023), 2% SDS (Invitrogen #15525017), 10% Glycerol (VWR #EM-GX0185-6)), run on a 10% polyacrylamide SDS gel at 115 V for 1.5-2 hr then transferred to 0.2 μm nitrocellulose membrane (Bio Rad #1620112) over night at 50 mA. Membranes were blocked with 5% milk in 1X TBS supplemented with 0.1% Tween-20 (1X TBST) for 1 hour then incubated for 2 hours or overnight in the indicated primary antibody. Membranes were rinsed 3x for 15 minutes in 1X TBST and incubated in the proper secondary antibody (Goat anti-mouse BioRad #170-6516 or Goat anti-rabbit BioRad #170-6515) for 1 hour. Membranes were rinsed 3x with 1X TBST for 15 minutes and developed using Clarity Western ECL Substrate (Bio-Rad, #1705060) and Molecular Imager Gel Doc (Bio Rad). The following antibodies were used for western blot analysis following the manufacturer’s instructions: N-Protein Antibody (SinoBiological, Rabbit mAb, 40068-RP02); HK2 Hexokinase II (HK2, Cell Signaling, Rabbit mAb, C64G5) Rabbit mAb Cell signaling; Hif1a (Novus Biologicals, Rabbit Ab, 102104-498); GAPDH (Cell Signaling, Rabbit mAb, 5174S); Glut-1 (Santa Cruz, Mouse mAb, # sc-377228); Viral proteins IE1, IE2^56^, pp28^57^, UL26^58^ (Mouse mAb).

For quantification of western blots GAPDH, HK2 and Glut1 (Fig. 4f), a single lysate sample of HFFs treated with vehicle or TNFα for 24 hr was run on three independent gels and immunoblotted for GAPDH, HK2 and Glut1 as described above. Band intensities were determined using Image Lab v4.1. Normalized GAPDH values were calculated by dividing each sample’s GAPDH intensity by the mean vehicle-treated GAPDH intensity of the three technical replicates. HK2 and Glut1 band intensities for each replicate were then divided by their respective normalized GAPDH values. Sample intensities were expressed as percentages of the highest band intensity (100) and averaged.

### Analysis of RNA

Analysis of RNA was carried out as previously described^52^. Briefly, RNA was extracted using Trizol (Invitrogen #15596026) and cDNA synthesis was completed using qScript cDNA Synthesis Kit (#95047-500). Expression of RNA was quantified by RT-qPCR for each indicated gene using the following primers: OC43: 5’- GGATTGTCGCCGACTTCTTA-3’ (forward) and 5’- CACACTTCTACGCCGAAACA-3’ (reverse). SARS-CoV-2: 5’-ATGAGCTTAGTCCTGTTG-3’ (forward) and 5’-CTCCCTTTGTTGTGTTGT-3’ (reverse). Human HK2 5’- GCCTACTTCTTCACGGAGCT-3’ (forward) and 5’-ATGAGACCAGGAAACTCTCG-3’ (reverse). Human Hif1α 5’-CGTTCCTTCGATCAGTTGTC-3’ (forward) and 5’- TCAGTGGTGGCAGTGGTAGT-3’ (reverse). Human Glut1: 5’- GCCTTCTTTGAAGTGGGTCC- 3’ (forward) and 5’- AGTTGGAGAAGCCTGCAACG-3’ (reverse). Gene expression was normalized to Human GAPDH: 5’-CATGTTCGTCATGGGTGTGAACCA-3’ (forward) and 5’- ATGGCATGGACTGTGGTCATGAGT-3’.

### CRISPR Knockout

CRISPR knockouts (KO) were performed with the Neon™ Transfection System 10 uL kit (ThermoFisher #MPK1025). HFFs were grown to 70% confluence and trypsinized using TrypLE Express (Invitrogen #12605010). HFFs were collected via centrifugation at 600 RPM for 5 minutes and resuspended to a concentration of 1.1×10^7^ in resuspension buffer R (ThermoFisher #MPK1025). In a separate tube, 60 pmol sgRNA was combined with 20 pmol Cas9 protein (Synthego) in a volume of 3 uL and incubated at room temperature for 15 minutes. To the prepared gRNA:Cas9 mixture, 9 uL cell solution (2×10^5^ HFFs) was added and gently pipetted up and down to mix. A 10 uL Neon™ pipette tip was used to extract 10 uL of the cell/gRNA/Cas9 solution which was then electroporated using the Neon™ Transfection System (voltage: 1650 V, width: 10 ms, pulses: 3) and transferred to a prepared 6-well dish (Greiner #82050-842) containing growth medium. This process was repeated with the same pipette tip and gRNA/Cas9 solution and transferred to the same dish for 2 transfections per guide into a single well. Guide RNAs were ordered from Synthego - Negative Control Scrambled sgRNA (modified) #1: GCACUACCAGAGCUAACUCA, HK2 (Gene Knockout Kit v2): sg1-CAUGCACGGCACCGGGGACG sg2-UCCGUGUUCGGAAUGGGAAG sg3-UCCAGAGAAAGGGGACUUCU, HIF1α (Gene Knockout Kit v2): sg1-AGGAAAGUCUUGCUAUCUAA sg2-UUCACAAAUCAGCACCAAGC sg3-ACACAGGUAUUGCACUGCAC.

### Synthego ICE score^59^

Knockout confirmation of Hif1α using Synthego’s Inference of CRISPR Edits (ICE). Genomic DNA was extracted from a 10 cm dish of sub-confluent HFFs transfected with Hif1α sgRNA/Cas9 (described above) or non-target guice control (ntg) sgRNA/Cas9 ribonucleoprotein complex using Lucigen quick extract DNA kit (item # QE09050). Hif1α gene locus surrounding the sgRNA target site was amplified from Hif1α edited DNA extract sample and ntg control DNA extract sample with Hif1α primer set 1 (sequences below) using touchdown PCR method. The resulting DNA fragment was PCR purified using Qiagen QIAquick PCR purification kit (item # 28104) and used as the DNA template in a second PCR method using Hif1α primer set 2 (Sequences below). The resulting DNA fragment was purified as described above and submitted to Genewiz for Sanger Sequencing using Hif1α primer set 3 (sequences below). Sequencing data for Hif1α edited and ntg control samples were uploaded to ice.synthego.com for analysis^59^. Primer sequences: Hif1α Set 1: F-GGGAAGGTTTACAGTTCCATGG; R-GTCTTGCTCTGTCATCCAGG. Hif1α Set 2: F- TCCAGGCTTAATCAGTTGGC; R-CTCAGCTCACCACAACATCC. Hif1α Set 3: F- GCAGCCTAGACTTTATACGAGG; R-ATCTCCTGACCTCAGATGATCC.

### Glucose Consumption

HFFs were grown to confluence in a 6-well TC-treated plate (Greiner #82050-842) and treated with 1 mL indicated treatment master mix for 24 hours. Medium was harvested from each well and glucose concentration for each medium sample was quantified using the HemoCue Glucose 201 System (HemoCue).

A standard curve was prepared using virgin medium where the highest dilution was 225 mg/dL glucose. This standard was serially diluted in PBS 1:2 five times for the remaining standards. To detect glucose, 8 μL each standard was loaded into a HemoCue glucose microcuvette (Hemocue #10842-830) and inserted into the glucose meter; mg/dL glucose detected was recorded and plot against the known concentration of glucose in the standards. Samples were diluted 1:4 In PBS and loaded into microcuvettes as described for standards.

To calculate glucose consumption, mg/dL glucose was determined for each sample using the standard curve then multiplied by its dilution. The amount of glucose detected in each sample was subtracted from the amount of glucose in t0 medium to quantify total nmol glucose consumed over 24 hr. The total amount of glucose consumed (nmol/hrx1e6 cells) was calculated by dividing the calculated nmol glucose consumed for each sample by 24 hr and the average HFF cell count in a 6-well dish (3.1e5 cells).

### Lactate Secretion

HFFs were grown to confluence in a 12-well TC-treated plate and treated with 500 μL indicated treatment master mix for 24 hours. Medium was harvested from each well and stored at -80C. To measure lactate in the medium, a standard curve was prepared using a 16 mM lactate standard as the most concentrated standard and serially diluting this most concentrated standard 1:2 8x. Medium samples were diluted 1:64 in PBS. Samples and standards were loaded onto the same MS run and resulting MS intensity data of the standards was used to generate a standard curve. Standards that began to plateau or fell below the limit of detection (3x the value of the average blank) were discarded from the standard curve. Sample MS intensity values that did not fall within the standard curve were discarded.

Lactate secretion for each sample was calculated by converting the MS intensity value to mM lactate using the standard curve and multiplying by the dilution factor (1:64) to yield lactate concentration (nmol/μL). Total nmol for each sample was calculated by multiplying the lactate concentration by the total sample volume (500 μL). Lactate secretion (nmol/hr*1e6 cells) was calculated by dividing the total nmol lactate secreted for each sample by 24 hr and the average HFF cell count in a 12 well plate (1.2e5 cells).

### Steady State Metabolomics Analysis

For steady state metabolomics, HFFs were grown to confluence in 10 cm TC-treated dishes (Greiner #82050-916) and placed in serum-free medium supplemented with 10 mM HEPES for 24 hours. HFFs were treated with 7 mL treatment medium supplemented with 10 mM HEPES for 24 hours. Medium was aspirated and metabolites were extracted from cells immediately by adding 3 mL cold 80% methanol (−80C). Extract was centrifuged at 3,000 RPM and supernatant containing metabolites was decanted into a fresh 50 mL conical on dry ice. Residual metabolites were extracted from the cell pellet with 3x washes of 500 uL 80% cold methanol followed by centrifugation at 3,000 RPM. Supernatants were pooled in the appropriate 50 mL conical and the methanol was evaporated under a gentile stream of nitrogen for 6-8 hours. Samples were suspended in 200 uL 80% cold methanol and analyzed by LC-MS/MS as previously described^60^ along 8x standards serially diluted 1:2 where the most concentrated standard contained: Glucose-6-phosphate (25 μM), Glucose-1-Phosphate (25 μM), Fructose-6-Phosphate (2.5 μM), Fructose-1,6-Bisphoaphate (20 μM), UDP-Glucose (50 μM), N-Acetyl Glucosamine-6-Phosphate (20 μM), UDP-N-Acetyl Glucosamine (40 μM) and N-Acetyl Glucosamine-1-Phosphate (20 μM). Absolute abundances of UDP-Glc and UDP-GlcNAc (Fig 6b) were calculated by converting MS intensity to nmol/μL UDP-Glc or UDP-GlcNAc using the appropriate standard curve. Total metabolite (nmol) in each sample was determined by multiplying total sample volume (200 μL) by the calculated metabolite concentration (nmol/μL). Total intracellular metabolite abundance (nmol/1e6 cells) was determined by dividing the nmol metabolite by the average HFF cell count in a 10 cm dish (1.7e6 cells).

For metabolomics data analysis, LC-MS/MS intensity values were imported to Metaboanalyst 5.061. Data was mean-centered and divided by the standard deviation (SD) of each variable. For principal component analysis (Fig 1a & Fig 3a), each dot represents metabolomics data of one biologically independent sample. Shaded ellipses represent 95% confidence intervals. PC1 and PC2 refer to the amount of total variation observed between samples that can be attributed to segregation along that principal component. Hierarchical clustering maps (Fig 1b & Fig 3b) follow Euclidian distance and Ward cluster algorithm. Metabolite values are depicted as Z-scores from min (blue) to max (red). Metabolites determined to be significantly different between TNFα and vehicle treatments (Fig 1c) were determined in Metaboanalyst. Log_2_ fold change (FC) values and FDR-adjusted p-values were calculated using Metaboanalyst. The direction of comparison was TNFα+/Vehicle. FC threshold of 1.5 was and FDR-adjusted p-value threshold of 0.05 were applied to yield 12 metabolites significantly altered as a result of TNFα treatment.

### Metabolic tracer analysis

HFFs were grown to confluence in 10 cm dishes in growth medium. Medium was replaced with serum-free medium containing 10 mM HEPES for 24 hours. Cells were then treated with 7 mL indicated treatment medium containing ^12^C-Glucose (4.5 g/L) and 10 mM HEPES for 19 hours. At 19 hours, medium was replaced with 7 mL indicated treatment medium containing U-^13^C-Glucose (Cambridge Isotopes #CLM-1396-1) (4.5 g/mL). Metabolites were extracted from cells as described above at 19 hours post treatment (t = 0 hours post label), 20 hours post treatment (t = 1 hours post label), 21 hours post treatment (t = 2 hours post label) and 25 hours post treatment (t = 5 hours post label). Samples were prepared and analyzed by LC-MS/MS as described above^60^. To analyze the intracellular abundance of isotopologues as represented in Fig. 6c & d, each isotopologue species was divided by the sum of all isotopologues detected and multiplied by the intracellular abundance of UDP-Glc or UDP-GlcNAc determined in Fig. 6b.

### Proteomics Experiment and Data Analysis

Cells were grown to confluence in 10 cm dishes (VWR, #82050-916) and placed in serum-free medium for 24 hours. Cells were treated in 7 mL indicated treatment medium at t0. At 24 hr post treatment, the monolayer was washed 3x with cold PBS. All subsequent steps were performed on ice and with chilled 4C PBS. Cells were scraped in 3 mL PBS and transferred to 15 mL conical tubes. Each dish was washed with an additional 3 mL of PBS and combined with its respective cell suspension. For each sample, cells were pelleted via centrifugation at 3,000 RPM for 5 minutes. Supernatant was discarded and the pellet was washed with 500 μL cold PBS and pelleted again at 3,000 RPM for 5 minutes. Supernatant was removed and samples were brought to the Mass Spectrometry Resource Laboratory (MSRL) at University of Rochester Medical Center. Protein extraction and S-trap Digest was performed by the MSRL. Samples were run on Oribtrap Fusion Lumos and analyzed using Data-Independent Acquisition (DIA) to yield relative protein abundance for 3,780 proteins, data shown in Supplementary Table 4.0. Data uploaded to MetaboAnalyst 5.0, mean-centered and divided by the standard deviation (SD) of each variable. For principal component analysis (Figure 5a) each dot represents proteomics data of one biologically independent sample. Shaded ellipses represent 95% confidence intervals. PC1 and PC2 refer to the amount of total variation observed between samples that can be attributed to segregation along that principal component.

To observe the differences in abundance of metabolic enzymes between vehicle and TNFα treated cells, a list of genes involved in metabolism was retrieved from the UniProt Database on August 31^st^, 2021 to yield 4,432 human genes involved in metabolism (Supplementary Table 4.1). This list of genes was scanned against our proteomics data set of 3,780 proteins using a custom code in Python v 3.7/PyCharm Community 12.1. The resulting list of intersecting genes (562 genes) were uploaded to MetaboAnalyst 5.0. Data was mean-centered and divided by the standard deviation (SD) of each variable. For principal component analysis (Supplementary Figure 1) each dot represents proteomics data of one biologically independent sample. Shaded ellipses represent 95% confidence intervals. PC1 and PC2 refer to the amount of total variation observed between samples that can be attributed to segregation along that principal component. Hierarchical clustering maps (Fig 4a) follow Euclidian distance and Ward cluster algorithm. Values are depicted as Z-scores from min (blue) to max (red). Proteins determined to be significantly more or less abundance as a result of TNFα-treatment relative to vehicle were determined in Metaboanalyst. Log2 fold change (FC) values and FDR-adjusted p-values were calculated using Metaboanalyst (Supplementary Table 4.1). The direction of comparison was TNFα+/Vehicle. FC threshold of 1.5 was and FDR-adjusted p-value threshold of 0.05 were applied to yield 40 proteins significantly altered as a result of TNFα treatment.

For the analysis a specific protein of interest, normalized protein abundance was calculated by dividing each sample relative protein abundance by the average of the vehicle-treated relative protein abundance (Fig. 4c & 6a). Statistics were performed in GraphPad Prism v9.1.0.

To generate a list of proteins more abundant in TNFα treatment compared to TNFα+2DG co-treatment, statistical parameters were applied to the data [Log_2_ Fold Change (TNFα+2DG/TNFα) < -0.6, p-value < 0.05] (Supplementary Table 4.2, 239 proteins). The resulting list of proteins which were uploaded to Gene Ontology (GO) Analysis to identify the biological process most impacted, represented by FDR-values (Fig. 5b). The list of proteins more abundant in TNFα treatment compared to TNFα+2DG co-treatment was also scanned against the GO-term [GO:0034097] “Involved in Cytokine Signaling” (54 proteins). These genes were scanned against databases of known glycosylated proteins^62, 63^ using a customized R-script to identify glycosylated proteins within this subset of proteins more abundant in TNFα treatment that are depleted during TNFα+2DG co-treatment (30 proteins glycosylated, 24 proteins not glycosylated. Missing values were replaced with the minimum detected MS intensity detected by that sample (Supplementary Table 4.2, yellow highlighted cells). Relative protein abundance values for each protein was determined by dividing the sample normalized protein abundance by the average relative protein abundance of vehicle-treated for that protein. The top 15 proteins most strongly induced by TNFα treatment, either glycosylated or not glycosylated, were graphed (Fig 5c).

Similarly, a list of proteins more abundant in TNFα+2DG co-treatment compared to TNFα treatment was generated by applying statistical parameters to the original proteomics list of 3,780 proteins [Log_2_ Fold Change (TNFα+2DG/TNFα) < 0.6, p-value < 0.05] (Supplementary Table 4.3, 120 proteins). The resulting list of proteins was uploaded to Gene Ontology (GO) Analysis to identify biological processed most impacted, represented by FDR-values (Supplementary Figure 3)

### Statistics

All statistical analysis were carried out using GraphPad Prism 9 unless otherwise indicated. MetaboAnalyst 5.0 was used to perform PCA analysis and hierarchical clustering from raw LC-MS/MS intensity data mean-centered and divided by the standard deviation (SD) of each variable.

---

**Supplementary Figure 1.**
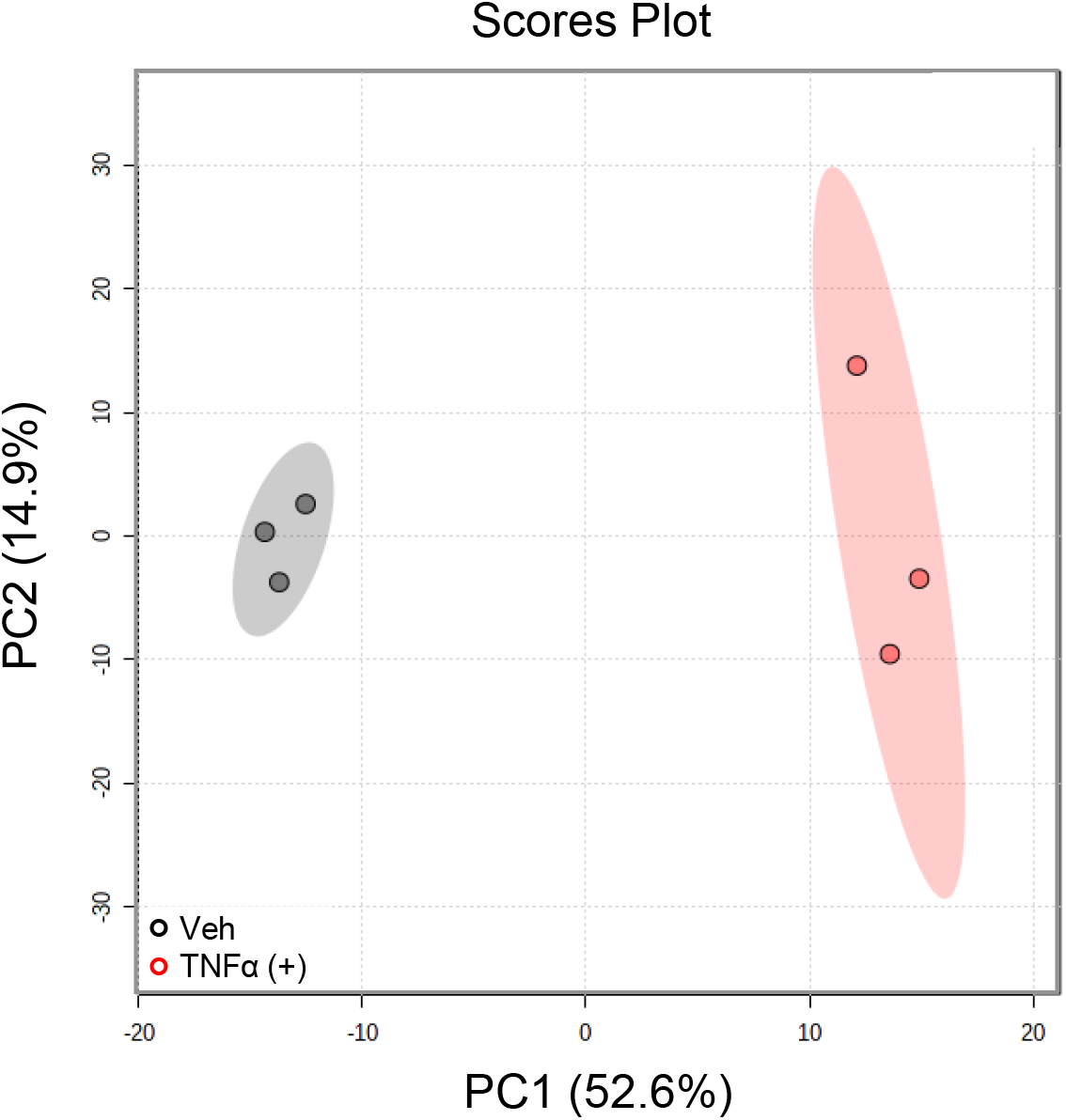
HFFs treated with vehicle (black) or TNFα (10 ng/mL, red) for 24 hr. Cells were harvested for proteomics analysis (Supplementary Table 4.0). PCA of a subset of proteins from proteomics analysis involved in metabolism; analysis described in materials and methods (Supplementary Table 4.1). Data shared with Fig. 4a.

**Supplementary Figure 2.**
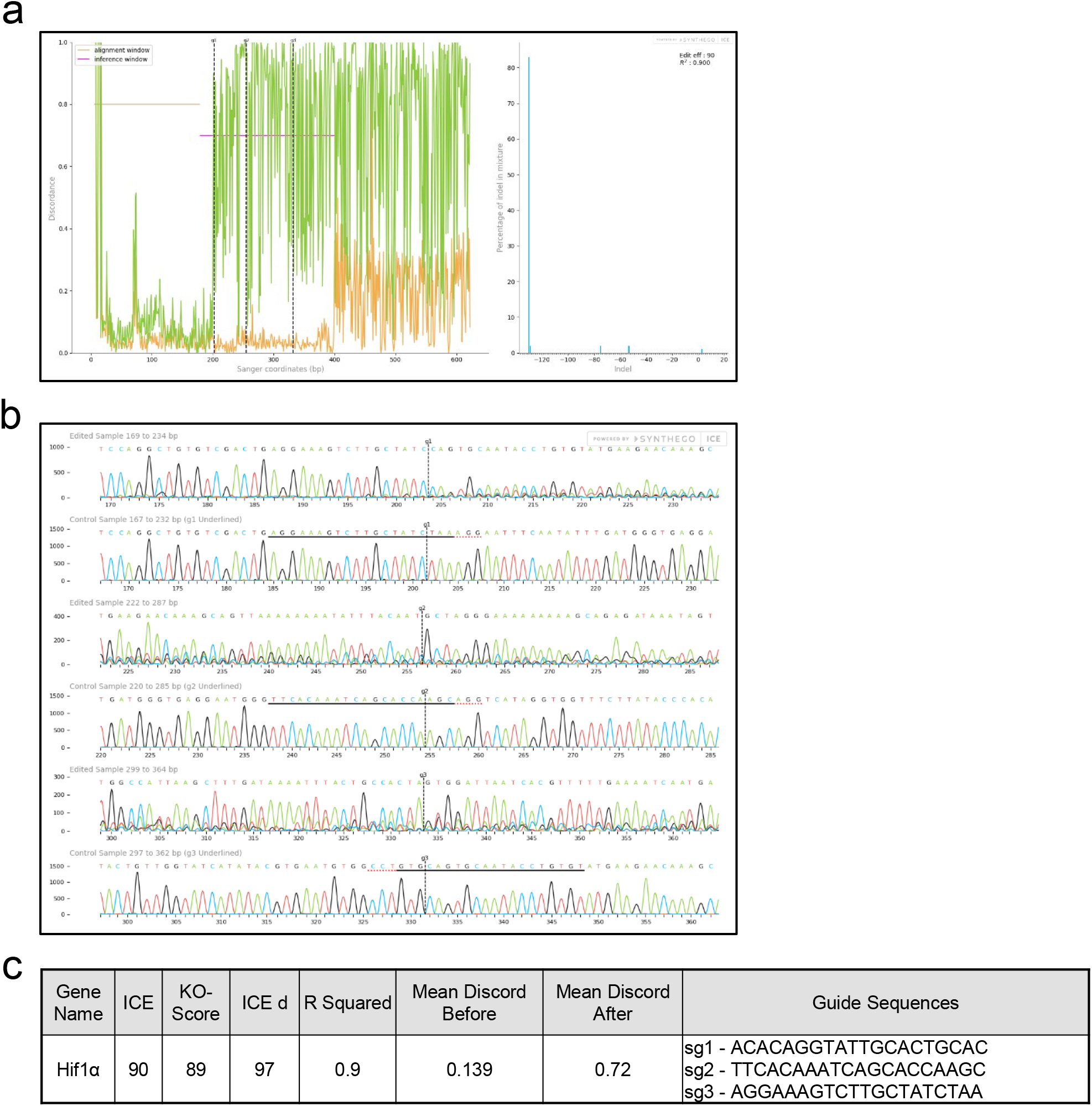
Knockout confirmation of Hif1α using Synthego’s Inference of CRISPR Edits (ICE) tool. HFFs treated with CRISPR Cas9-RNP containing guides for Hif1α or a non-target guide (ntg) to generate knockout (KO) cell lines. Hif1α and ntg targeted cells harvested for genomic DNA and HIF1A gene locus amplified with sequencing primers. Samples submitted for Sanger sequencing and results uploaded to Synthego’s ICE tool. **a** Alignment plot, left, showing control (orange) and edited (green) sequences. Vertical dotted lines indicate guide sequences in relation to Sanger sequence coordinates. Indel plot, right, showing the predicted range of insertions and deletions in the edited gene locus. **b** Trace files of Hif1α (edited sample) and ntg (control sample) targeted cells spanning the cut site of HIF1A gene locus targeted by sgRNAs. Guide sequences underlined by black solid line in the control trace, PAM sequences denoted by dotted red underline and vertical dotted lines indicate expected cut site. **c** Table from ICE analysis displaying ICE and KO score as well as sgRNA sequences towards HIF1A.

**Supplementary Figure 3.**
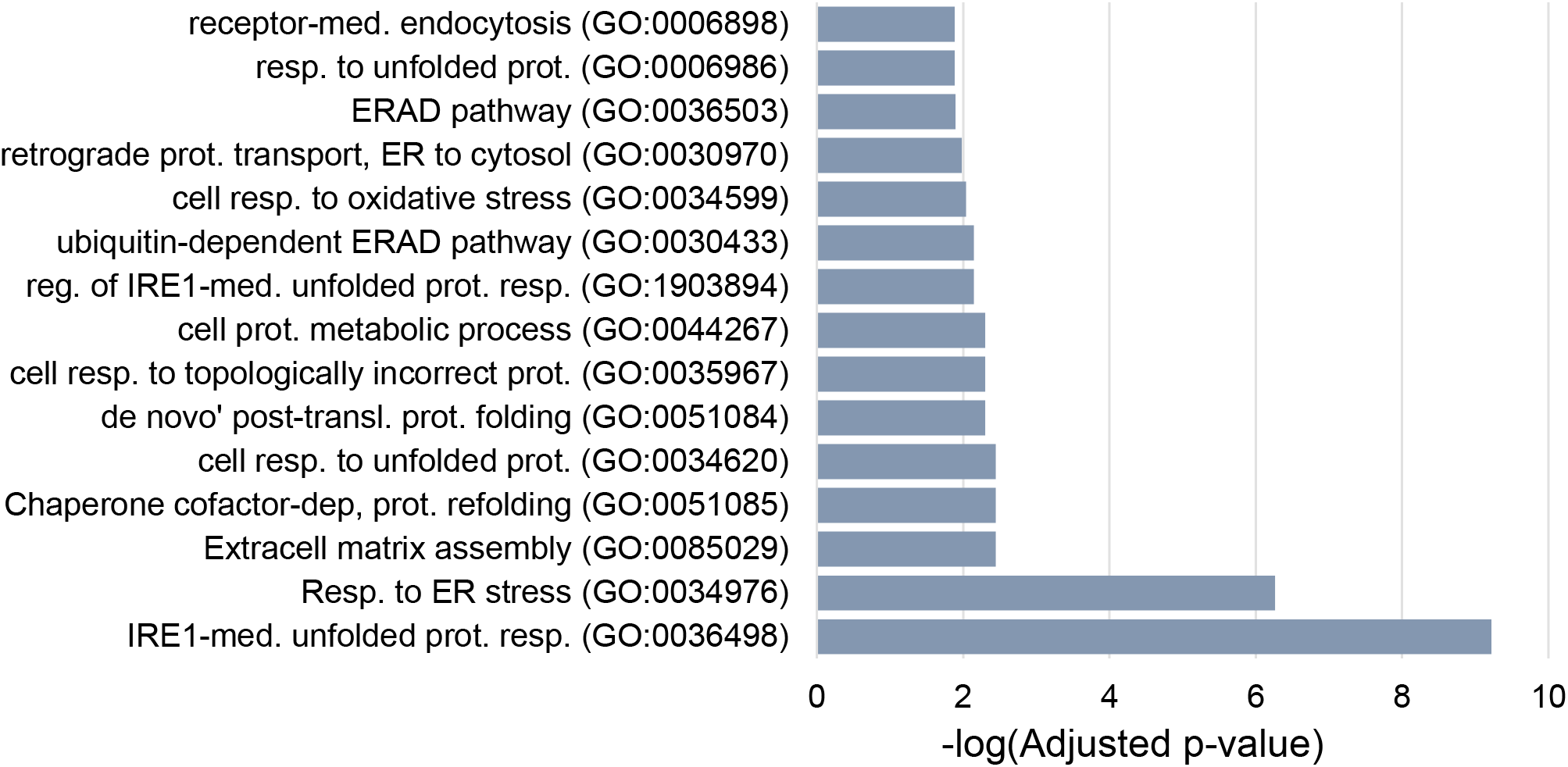
HFFs treated with vehicle and TNFα (10 ng/mL) in the presence and absence of 2DG (20 mM) for 24 hr. Cells harvested and analyzed for protein abundance (Supplementary Table 4). Ontology analysis of TNFα-induced proteins that were significantly more abundant upon co-treatment with 2DG. Bar graph represents the FDR-values from the top 15 GO-terms (Supplementary Table 4.3).

**Supplementary Figure 4.**
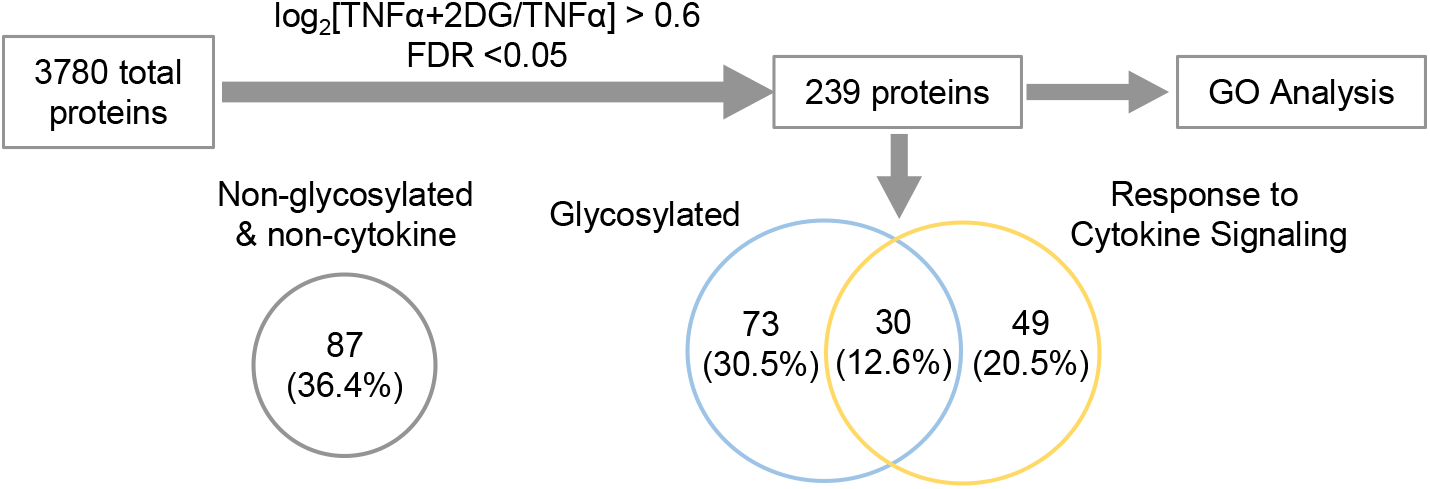
Proteomics analysis workflow. HFFs treated with vehicle and TNFα (10 ng/mL) in the presence and absence of 2DG (20 mM) 24 hr. Cells harvested and analyzed for protein abundance (Supplementary Table 4). Statistical parameters applied to proteomics data, described in materials and methods, to generate a list of proteins significantly induced by TNFα treatment but depleted in cells co-treated with TNFα and 2DG. Protein list submitted for Gene Ontology (GO) analysis (Fig. 6b) and scanned against databases of known glycosylated proteins (described in materials and methods) or ‘Response to Cytokine Signaling’ [GO:0034097] (Supplementary Table 4.2). Venn Diagram represents overlap of proteins involved in cytokine signaling (yellow) and glycosylated (blue).

